# Can Actin Depolymerization Actually Result in Increased Plant Resistance to Pathogens?

**DOI:** 10.1101/278986

**Authors:** Hana Krutinová, Lucie Trdá, Tetiana Kalachova, Lucie Lamparová, Romana Pospíchalová, Petre I. Dobrev, Kateřina Malínská, Lenka Burketová, Olga Valentová, Martin Janda

## Abstract

The integrity of the actin cytoskeleton is essential for plant immune signalling^1^. Consequently, it is generally assumed that actin disruption reduces plant resistance to pathogen attack^2^-^4^. However, in a previous study, it was shown that actin depolymerisation triggers the salicylic acid (SA) signalling pathway^5^, which is interesting because increased SA is associated with enhanced plant resistance to pathogen attack^6^,^7^. Here, we attempt to resolve this seeming inconsistency by showing that the relationship between actin depolymerization and plant resistance is more complex than currently thought. We investigate the precise nature of this relationship using two completely different plant pathosystems: i) a model plant (*Arabidopsis thaliana*) and a bacterial pathogen (*Pseudomonas syringae*), and ii) an important crop (*Brassica napus*) and a fungal pathogen (*Leptosphaeria maculans*). We demonstrate that actin depolymerization induces a dramatic increase in SA levels and that the increased SA is biosynthesized by the isochorismate synthase pathway. In both pathosystems, this phenomenon leads to increased plant resistance.

## Main text

The actin cytoskeleton plays a key role in plant immunity^1^, both by providing a physical barrier and by its involvement in the transport of callose, antimicrobial compounds and cell wall components to an infection site^8^. Several studies have shown that when drugs, such as cytochalasin E or latrunculin B, depolymerize the actin cytoskeleton, different plant species become more susceptible to pathogens. For example, treatment of *A. thaliana* with latrunculin B resulted in higher susceptibility to infection by *Pseudomonas syringae*^3^,^9^,^10^. Also it was shown that *P. syringae* secrets HopW1 effector which disrupts actin cytoskeleton^9^,^10^. Furthermore, treatment with cytochalasin E increased the penetration of *A. thaliana* plants by *Colletotrichum* species^4^ and the rate of entry to barley by *Blumeria graminis* f. sp. *hordei* ^2^. However, in tobacco, cytochalasin E induced the transcription of *NtPR-1* (pathogenesis-related 1), a defence-related SA marker gene^11^. Furthermore, both cytochalasin E and latrunculin B induced the transcription of several SA marker genes (*AtPR-1, AtPR-2* and *AtWRKY38*) in *A. thaliana* seedlings^5^. This suggests that while such drugs do indeed cause actin depolymerization, the effects of such depolymerization may not always be adverse. Could it be that drug-induced actin depolymerization actually triggers processes that induce the SA pathway and thereby increase plant resistance to pathogens?

To establish that SA levels can increase upon actin depolymerization, we measured phytohormone content in *A. thaliana* seedlings treated with just 200 nM latrunculin B. Such a low concentration of latrunculin B proved sufficient to depolymerize actin filaments in the seedlings within 24 h (Fig. S1). Significantly, by that time there was a sevenfold increase in the free SA level of the treated seedlings compared with the control ones. The only other phytohormone to display an increase (twofold) was jasmonic acid (JA). Apart from Indole-3-acetamide (IAM), which showed a threefold decrease, the other tested phytohormones remained largely unaltered (Fig. 1A; Table S1).

**Fig. 1.**
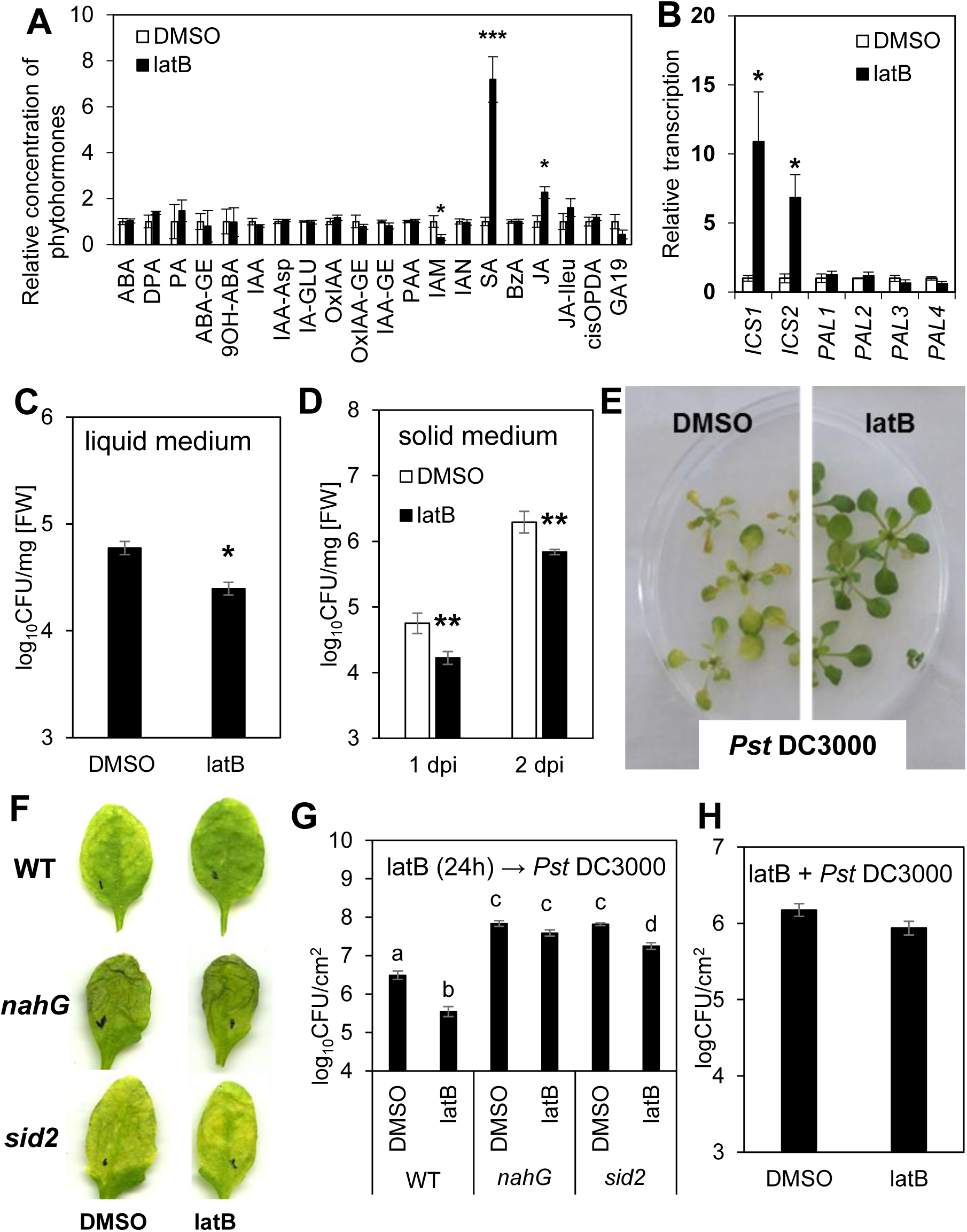
Effects of latrunculin B on *Arabidopsis thaliana*. **A)** Phytohormone analysis of seedlings grown *in vitro* in liquid MS medium. Seedlings were treated for 24 h with 200 nM latrunculin B (latB) or 0.01% DMSO (control). For abbreviations of analyzed phytohormones, see Table S1. **B)** Transcription of SA biosynthetic genes *ICS1, ICS2, PAL1, PAL2, PAL3* and *PAL4.* Seedlings were treated for 24 h with 200 nM latB or 0.01% DMSO. The transcription level was normalized to the reference gene, *SAND*. **C)** Bacterial titres in seedlings grown in liquid MS medium. Seedlings were pretreated for 24 h with 200 nM latB or 0.01% DMSO before inoculation with *Pst* DC3000. Tissue was harvested 1 day after inoculation with bacteria. **D)** Bacterial titres in seedlings grown in solid MS/2 medium. Seedlings were pretreated for 24 h with 200 nM latB or 0.01% DMSO before inoculation with *Pst* DC3000. Tissue was harvested 1 and 2 days after inoculation with bacteria. **E)** Representative photographs of seedlings grown on MS/2 medium 2 days after inoculation with *Pst* DC3000. **F)** Representative photographs of adult *A. thaliana* leaves infected with *Pst* DC3000 3 days after inoculation. **G,H)** Bacterial titres in adult plants. **G)** Plants were treated for 24 h with 1 µM latB or 0.05% DMSO before inoculation with *Pst* DC3000. **H)** Plants were treated with 1 µM latB or 0.05% DMSO, each in a solution containing *Pst* DC3000. Tissue was harvested 3 days after inoculation with *Pst* DC3000. *A. thaliana* WT plants (col-0) and mutants with impaired SA pathways (*nahG* and *sid2*) were used (G). The values represent mean and error bars (SEM) from four (A, C), three to four (B), five (D), and six (G, H) independent samples. The asterisks represent statistically significant changes in latB-treated samples compared with controls (* P<0.05; ** P<0.01; *** P<0.001; two tailed Student’s t-test).

Having shown this dramatic rise in SA level in *A. thaliana*, we wondered which of its two SA biosynthetic pathways was responsible for this increase or whether they both contributed to it. One pathway involves phenylalanine ammonia-lyase (PAL, EC 4.3.1.24), which exists in four isoforms, while the other involves isochorismate synthase (ICS; EC 5.4.4.2), which occurs in two isoforms^12^. Analysis of the transcription of all *AtPAL* and *AtICS* genes in the seedlings revealed that only the *AtICS* genes were induced by latruculin B (Fig. 1B). This shows that drug-induced actin depolymerization activates the ICS-dependent pathway and that this pathway alone is responsible for SA biosynthesis under these conditions.

Given that increased resistance to pathogens in *A. thaliana* is associated with SA biosynthesis through the ICS pathway^13^,^14^, is it possible that activation of the same pathway invoked by drug-induced actin depolymerization also results in increased resistance? To investigate this, we used Ishiga et al.‘s (2011) protocol as a basis for performing two *in vitro A. thaliana-Pseudomonas syringae* pv. *tomato* DC3000 (*Pst* DC3000) flood-inoculation assays in liquid and solid media^15^. We treated the seedlings with latrunculin B 24 h before inoculation with *Pst* DC3000. Remarkably, under both conditions, the latrunculin B-pretreated seedlings were more resistant than the pretreated control ones (Fig. 1C, D, E). To ensure that this phenomenon is not just associated with *in vitro* conditions, we also performed experiments using four-week old *A. thaliana* plants cultivated in soil, such plants typically being used for studies of *A. thaliana* resistance to *Pst* DC3000^16^. Unlike the seedlings, treatment with 200 nM latrunculin B did not activate the SA pathway in the adult plants and, thus, no increased resistance was observed (Fig. S2A, C). However, the transcription of SA marker genes (*AtPR-1, AtPR-2, AtICS1*) was induced after treatment with 1 µM latrunculin B (Fig. S2B), leading to increased resistance to *Pst* DC3000 (Fig. 1F, G, S2C). This suggests that plant resistance is strongly dependent on latrunculin B concentration, probably due to differences between the efficiency of latrunculin B-induced actin depolymerization in seedlings and in adult plants (Fig. S3). Similar to latrunculin B, pretreatment with cytochalasin E led to both SA-induced gene transcription (Fig. S4A) and increased plant resistance to *Pst* DC3000 (Fig. S4B), thereby strengthening the notion that such resistance is due to the depolymerizing activity of cytoskeletal drugs. It should be noted that we exclude the antibacterial effect of latrunculin B because *Pst* DC3000 grew *in vitro* in the presence of latrunculin B at a similar rate as in the control medium (Fig. S5A).

To further demonstrate the dependence of such resistance on the SA pathway, we performed assays using mutants known to have an impaired SA pathway and thus be more susceptible to *Pst* DC3000: *nahG*, which induces low endogenous SA levels through the expression of SA-hydroxylase, and *sid2*, a knock-out mutant of the *AtICS1* gene. As expected, latrunculin B did not induce resistance in the *nahG* plants (Fig. 1F, G). The *sid2* plants exhibited latrunculin B-induced resistance, but to a much lesser extent than the WT controls (Fig. 1F, G); this might be explained by induced transcription of the *AtICS2* gene after treatment with latrunculin B (Fig. 1B). Both of these results confirm that SA is indeed responsible for actin depolymerization-induced resistance.

To show that this phenomenon is not species-specific, we investigated the effect of latrunculin B on another model plant, *Nicotiana bentamiana*, and an important crop, oilseed rape (*Brassica napus*). As in the case of *A. thaliana*, treatment with 200 nM latrunculin B for 24 h induced the transcription of SA marker genes (*NbPR-1, NbPR-2*) in 14-day-old *N. benthamiana* seedlings *in vitro* (Fig. 2A). Likewise in *B. napus*, latrunculin B upregulated the transcription of SA marker genes (*BnPR-1, BnICS1*) (Fig. 2B). Furthermore, as with adult *A. thaliana*, the effect of latrunculin B on *B. napus* was concentration dependent (Fig. 2A). The treatment of *B. napus* with 10 µM latrunculin B 3 days before inoculation with a hemibiotrophic fungal pathogen, *Leptosphaeria maculans*, efficiently inhibited hyphal colonisation and necrosis formation in the infected cotyledons (Fig. 2C, E, F). Treatment with 1 µM latrunculin B led to weak and variable resistance against *L. maculans* (Fig. S6), corresponding to the weaker transcription of defence-related genes (Fig. 2B). These data are in accordance with our previous study characterizing the importance of SA in the defence of *B. napus* against *L. maculans*^17^. In addition, we observed significant cytochalasin E-induced resistance to *L. maculans* in *B. napus* (Fig. S7), which suggests that the effect is not compound-specific. Furthermore, neither latrunculin B nor cytochalasin E displayed antifungal activity on *L. maculans* growth *in vitro* (Fig. S5). Interestingly, the co-inoculation of *B. napus* cotyledons with a joint solution of 10 µM latrunculin B and *L. maculans* conidia also induced resistance (Fig. 2D). These results indicate that depolymerized actin can trigger resistance to bacterial or fungal pathogens.

**Fig. 2.**
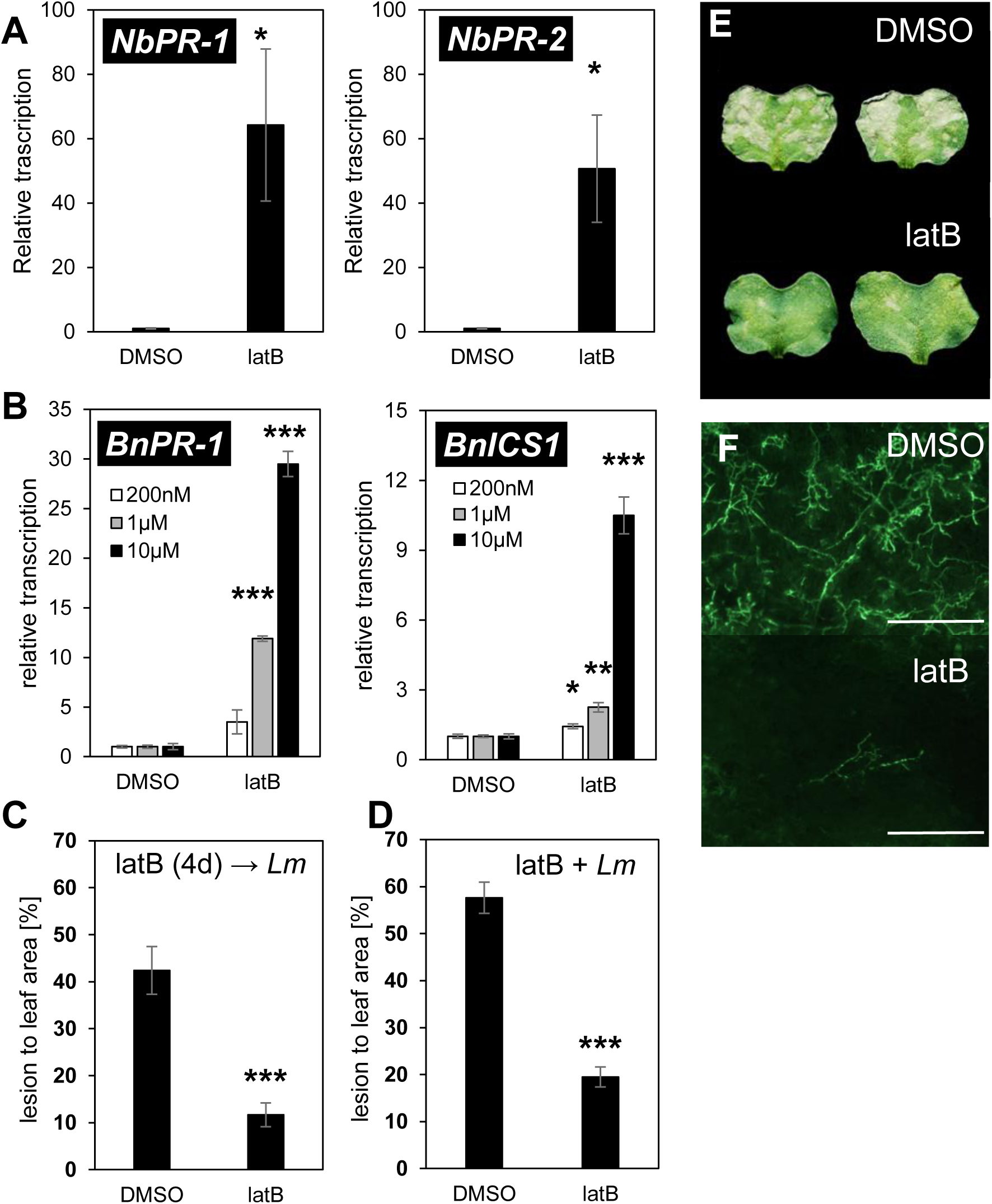
Effects of latrunculin B on *Nicotiana benthamiana* and *Brassica napus*. **A)** Transcription level of SA marker genes *NbPR-1* and *NbPR-2* in two-week-old *N. benthamiana* seedlings. Seedlings were treated for 24 h with 200 nM latB. Control cotyledons were treated for 24 h with a corresponding concentration of DMSO (0.01%). The transcription level was normalized to the reference gene, *NbSAND*. **B)** Transcription of SA marker genes *BnPR-1* and *BnICS1* in *B. napus* cotyledons. Cotyledons were treated for 24 h with infiltrations of 0.2, 1 or 10 µM latrunculin B (latB). Control cotyledons were treated for 24 h with a corresponding concentration of DMSO (0.01, 0.05 or 0.5%). The transcription level was normalized to the reference gene, *BnTIP41*. **C, D)** *B. napus* susceptibility to *L. maculans* was evaluated as the relative lesion area (ratio of lesion area to whole leaf area) on the cotyledons. Cotyledons treated with 10 µM latB 4 days before inoculation with *L. maculans* (**C**); cotyledons inoculated with a joint solution containing both 10 µM latB and *L. maculans* (**D**). Control cotyledons were treated with 0.5% DMSO. **E)** Representative images of *L. maculans*-infected cotyledons. **F)** Representative microscopy images of *L. maculans* hyphae proliferation in *B. napus* cotyledons in response to 10 µM latB or 0.5% DMSO. The bars correspond to 500 µM. The values represent mean and error bars (SEM) from three to four (A, B) and 18-24 (C, D) independent samples. The asterisks represent statistically significant changes in latB-treated samples compared with controls (* P<0.05; ** P<0.01; *** P<0.001; two tailed Student’s t-test).

Thus, we have shown that plant immunity is strongly activated by depolymerised actin and that this phenomenon appears to be generally valid; namely, it seems not to be species specific, pathogen-type specific or drug-type specific. These findings do not negate those of previous studies that showed the susceptibility of plants treated with cytoskeletal drugs to pathogens^2^-^4^,^9^,^10^. Rather, they reveal that the plant disease resistance is strongly dependent on whether the plant has sufficient time to activate SA-mediated immunity. This was clearly shown by our experiments with *Pst* DC3000, in which pre-infection treatment with cytoskeletal drugs resulted in resistance whilst co-inoculation did not (Fig. 1G, H). Interestingly, treatment with latrunculin B resulted in increased resistance in both *L. maculans* setups: pretreatment (Fig. 2C) and co-inoculation (Fig. 2D). This suggests that the rapidity of pathogen growth is a crucial factor. In contrast to *Pst* DC3000, which strongly damaged the inoculated leaves within three days, almost no multiplication of *L. maculans* occurred during the same period^17^. Thus, it appears that the slow growth of *L. maculans* enabled *B. napus* to establish the SA pathway, which was induced within 24 hours of cytoskeletal drug treatment (Fig. 2B). Overall then, while it is true that plant resistance to pathogens is decreased by a disrupted actin cytoskeleton, our results show that, given sufficient time, plants are able to trigger SA-based defence mechanisms to overcome such threats.

To our best knowledge, we herein provide the first evidence that disruption of the actin cytoskeleton can actually lead to increased plant resistance to pathogens, and that SA is crucial to this process. We strongly believe that our work opens a new and important direction for further research. For example, it is possible that plants have evolved a system for detecting actin cytoskeleton disruption and that the activation of such a system triggers SA-specific immune responses. For this reason, further research should be focused on deciphering the mechanism by which actin depolymerization triggers SA biosynthesis.

## MATERIAL AND METHODS

### Plant material

For the *Arabidopsis thaliana* experiments, the following genotypes were used: Columbia-0 (WT); *sid2-3* (SALK_042603)^18^; *nahG*^6^; *pUBC*::Lifeact-GFP^19^ and *p35S*::GFP-FABD2^20^. Arabidopsis seedlings were grown either in liquid MS medium or on solid MS/2 medium. Per litre, the liquid MS medium contained the following: 4.41 g Murashige and Skoog medium including vitamins (Duchefa, Netherlands), 5 g sucrose, 5 g MES monohydrate (Duchefa, Netherlands). Per litre, the solid MS/2 medium contained 2.2 g Murashige and Skoog medium (Duchefa, Netherlands) with 10 g sucrose and 8 g Plant agar (Duchefa, Netherlands). Both media were adjusted to pH 5.7 using 1M KOH. For cultivation in the liquid, surface-sterilized seeds were sown in 24-well plates containing 400 μL of liquid MS medium per well. The plants were cultivated for 10 days under a short-day photoperiod (10 h/14 h light/dark regime) at 100-130 μE m^-2^ s^-1^ and 22 C. On the 7th day, the medium in the wells was exchanged for a fresh one. For cultivation on the solid MS/2 medium, seedlings were grown in Petri dishes for 12 days under a long-day photoperiod (16 h/8 h light/dark regime) at 100-130 μE m^-2^ s^-1^ and 22 °C. For *A. thaliana* plants grown for 4 weeks in soil, surface-sterilized seeds were sown in Jiffy 7 peat pellets and the plants cultivated under a short-day photoperiod (10 h/14 h light/dark regime) at 100-130 μE m^-2^ s^-1^, 22 °C and 70% relative humidity. They were watered with fertilizer-free distilled water as necessary

For the *Brassica napus* experiments, plants of the Eurol cultivar were grown hydroponically in perlite in Steiner’s nutrient solution^21^ under a 14 h/10 h light/dark regime (25 °C/22 °C) at 150 μE m^−2^ s^−1^ and 30-50% relative humidity. True leaves were removed from 14-day-old plantlets to avoid cotyledon senescence.

For the *Nicotiana benthamiana* experiments, surface-sterilized seeds were grown in 24-well plates containing 400 μL of liquid MS medium for 14 days under a 10 h/14 h light/dark regime at 100-130 μE m^-2^ s^-1^ and 22 °C. On the 7th day, the medium in the wells was exchanged for a fresh one.

### Treatment with chemical compounds

As actin depolymerizing drugs, latrunculin B (Sigma-Aldrich, USA) and cytochalasin E (Sigma-Aldrich, USA) were used. Latrunculin B and cytochalasin E were both dissolved in DMSO; the concentration of the stock solutions were 2 mM and 4 mM, respectively.

For the *Pst* DC3000 resistance assay, the seedlings grown in 24-well plates were treated by replacing the pure liquid MS medium in the plate wells with medium containing 200 nM latrunculin B or 0.01% DMSO control. The seedlings cultivated on the solid medium were treated by flooding with 10 mL of MS/2 medium containing 200 nM latrunculin B or 0.01% DMSO control.

For the transcriptomic assay, the seedlings *of Arabidopsis thaliana* grown in 24-well plates were treated with 200 nM latrunculin B (0.01% DMSO control) or 10 µM cytochalasin E (0.25% DMSO control). The seedlings of *Nicotiana bentamiana* were treated with 200 nM latrunculin B (0.01% DMSO control).

Fully-developed leaves from four-week-old *A. thaliana* grown in soil were infiltrated either with 200 nM or 1 µM latrunculin B (0.01% or 0.05% DMSO as respective controls) or with 1 µM or 10 µM cytochalasin E (0.025% or 0.25% DMSO as respective controls) using a needleless syringe.

The 10-day-old cotyledons of *B. napus* were infiltrated either with 1 µM or 10 µM latrunculin B or with 10 µM cytochalasin E (in all cases with corresponding DMSO controls) using a needleless syringe.

### Inoculation of *A. thaliana* seedlings with *Pst* DC3000

After the *A. thaliana* seedlings had been cultivated in 24-well plates in the liquid MS medium for 10 days, the cultivation medium was exchanged for one containing latrunculin B or cytochalasin E, and incubated for 24 h. On day 11, the medium was replaced with a bacterial suspension of *Pst* DC3000 in 10 mM MgCl_2_ (OD_600_=0.01). The seedlings were incubated in this bacterial suspension for 1 min. After incubation, the suspension was replaced with the liquid MS medium. On day 12, the seedlings were harvested, each sample taken containing all of the seedlings from three wells. The seedlings were then homogenized in tubes with 1 g of 1.3 mm silica beads using a FastPrep-24 instrument (MP Biomedicals, USA). The resulting homogenate was serially diluted and pipetted onto King B plates. The colonies were counted after 1-2 days of incubation at 28 °C.

The seedlings cultivated on solid medium were flooded with 200 nM latrunculin B solution in water on day 13. Control plants were treated with a corresponding solution of DMSO. On day 14, the solutions were replaced with a suspension of overnight culture of *Pst* DC3000 (OD_600_=0.01) containing 0.025% Silwet. Samples were harvested at 0, 1 and 2 dpi, with each sample containing the plants from five plates. The seedlings were homogenized in tubes with 1 g of 1.3 mm silica beads using a FastPrep-24 instrument (MP Biomedicals, USA). The resulting homogenate was serially diluted and pipetted onto LB plates containing rifampicin. The colonies were counted after 1-2 days of incubation at 28 °C.

### Inoculation of four-week-old *A. thaliana* with *Pst* DC3000

*Pst* DC3000 was grown overnight on King B agar plates at 28 °C, resuspended in 10 mM MgCl_2_, and diluted to an OD_600_ of 0.001. Using a needleless syringe, the bacterial suspension was infiltrated into three fully-developed leaves from one plant. After 3 days, the infected tissue was collected as cut leaf discs (one disc per leaf, 0.6-cm diameter); three leaf discs from one plant represent one sample. The discs were homogenized in tubes with 1 g of 1.3 mm silica beads using a FastPrep-24 instrument (MP Biomedicals, USA). The resulting homogenate was serially diluted and pipetted onto King B plates. The colonies were counted after 1-2 days of incubation at 28 °C.

### Inoculation of *B. napus* with *Leptosphaeria maculans*

*L. maculans* isolate v23.1.3^17^,^22^ was used to inoculate *B. napus*. After harvesting, conidia obtained according to Šašek et al. (2012) ^17^ were washed once with distilled water, diluted to 10^8^ spores/ml, and stored at –20 °C for up to 6 months. The cotyledons of 14-day-old plants were infiltrated by conidial suspension (10^5^ conidia/ml), with at least 12 plants being used for each inoculation. The leaves were assessed for lesions 10 days after inoculation. The leaf area and the lesion areas therein were measured by image analysis using APS Assess 2.0 software (American Phytopathological Society, USA). The relative lesion area was then calculated as the ratio of lesion area to whole leaf area. For the microscopy studies, the cotyledons infected with GFP-tagged v23.1.3 isolate^23^ were observed at 10 dpi using a Leica DM5000 B microscope.

### Gene expression analysis

The whole seedlings from three independent wells were immediately frozen in liquid nitrogen. The tissue was homogenized in tubes with 1 g of 1.3 mm silica beads using a FastPrep-24 instrument (MP Biomedicals, USA). Total RNA was isolated using a Spectrum Plant Total RNA kit (Sigma-Aldrich, USA) and treated with a DNA-free kit (Ambion, USA). Subsequently, 1 μg of RNA was converted into cDNA with M-MLV RNase H^−^ Point Mutant reverse transcriptase (Promega Corp., USA) and an anchored oligo dT21 primer (Metabion, Germany). Gene expression was quantified by q-PCR using a LightCycler 480 SYBR Green I Master kit and LightCycler 480 (Roche, Switzerland). The PCR conditions were 95 °C for 10 min followed by 45 cycles of 95 °C for 10 s, 55 °C for 20 s, and 72 °C for 20 s. Melting curve analysis was then conducted. Relative expression was normalized to the housekeeping genes *AtSAND, AtTIP41, BnTIP41* and *NbSAND*. Primers were designed using PerlPrimer v1.1.21^24^. A list of the analysed genes and primers is available in Table S2.

### Phytohormonal analysis

Hormone analysis was carried out on four samples, each of which contained all seedlings from six of the 24 wells. Plant hormone levels were determined as described by Dobrev and Kaminek (2002)^25^. Briefly, samples were homogenized in tubes with 1.3 mm silica beads using a FastPrep-24 instrument (MP Biomedicals, USA). The samples were then extracted with a methanol/H_2_O/formic acid (15:4:1, v:v:v) mixture, which was supplemented with stable isotope-labeled phytohormone internal standards (10 pmol per sample) in order to check recovery during purification and validate the quantification. The clarified supernatants were subjected to solid phase extraction using Oasis MCX cartridges (Waters Co., USA). The eluates were evaporated to dryness and the generated solids dissolved in 30 μl of 15% (v/v) acetonitrile in water. Quantification was performed on an Ultimate 3000 high-performance liquid chromatograph (Dionex, USA) coupled to a 3200 Q TRAP hybrid triple quadrupole/linear ion trap mass spectrometer (Applied Biosystems, USA) as described by Dobrev et al. (2017)^26^. Metabolite levels were expressed in pmol/g fresh weight (FW).

### Confocal microscopy of actin filaments

For *in vivo* microscopy, a Zeiss LSM 880 inverted confocal laser scanning microscope (Carl Zeiss AG, Germany) was used with either a 40× C-Apochromat objective (NA = 1.2 W) or a 20x Plan-Apochromat objective (NA = 0.8). GFP fluorescence (excitation 488 nm, emission 489–540 nm) was acquired in z-stacks (20-25 µm thickness). The maximum intensity projections obtained from the z-stacks were created using Zeiss ZEN Black software.

### Growth of *Pst* DC3000 and *L. maculans in vitro* in presence of latrunculin B or cytochalasin E

*Pst* DC3000 grew overnight on solid LB medium containing rifampicin. From this, a fresh bacterial suspension was prepared (OD_600_=0.01) in liquid LB or liquid MS medium. To this suspension, latrunculin B (200 nM or 1 µM) or DMSO (0.05% or 0.01%) was added. The OD_600_ was measured 6 and 24 h after suspension preparation. Four independent samples were prepared for each type of treatment.

Conidia of the GFP-tagged v23.1.3 isolate of *L. maculans*^23^ were grown *in vitro* in Gamborg B5 medium (Duchefa, Netherlands) supplemented with 0.3% (w/v) sucrose and 10 mM MES monohydrate, and adjusted to pH 6.8. This medium contained latrunculin B (0.2, 1, 10 µM), cytochalasin E (1, 10 µM) or DMSO control (0.5%), and had a final concentration of 2500 conidia per well. The plates (black 96-well plate, Nunc R), covered with lids and sealed with Parafilm®, were incubated in darkness at 26 °C. On day 4, fluorescence was measured using a Tecan F200 fluorescence reader (Tecan, Switzerland) equipped with a 485/20 nm excitation filter and 535/25 nm emission filter. Eight wells were measured for each treatment.

### Statistical Analyses

All experiments were repeated at least three times. All statistical analyses were performed with Microsoft Excel 2013. The P values were calculated using a two-tailed Student’s t-test.

## Acknowledgement

We would like to thank Dr. Kenichi Tsuda from MPIPZ Cologne for providing the strain of *Pseudomonas syringae* pv. *tomato* DC3000. This work was supported by Czech Science Foundation grant no. 17-5151S and GAUK no. 992416. IEB Imaging Facility is supported by OPPK CZ.2.16/3.1.00/21519 and MEYS LM2015062. The pUBC::Lifeact-GFP and 35S::GFP-FABD2 seeds were kindly provided by Dr. Denisa Oulehlová from Žárský laboratory at IEB ASCR.

## Author contribution

H.K., L.T., T.K., M.J. designed the experiments; H.K., L.T., T.K., L.L., R.P., P.I.D., K.M., M.J. performed experiments; H.K., L.T., T.K., L.B., O.V., M.J. analysed the data; H.K., M.J. wrote the paper. All the authors discussed the results and commented on the manuscript.

**Correspondence and requests for materials** should be addressed to M. Janda.

